# Structural basis for FLCN RagC GAP activation in TFEB substrate-selective mTORC1 regulation

**DOI:** 10.1101/2022.06.28.498039

**Authors:** Rachel M. Jansen, Roberta Peruzzo, Simon A. Fromm, Adam L. Yokom, Roberto Zoncu, James H. Hurley

**Affiliations:** Department of Molecular and Cell Biology, University of California Berkeley; Berkeley CA 94720, USA; California Institute for Quantitative Biosciences, University of California, Berkeley, CA, 94720, USA; Helen Wills Neuroscience Institute, University of California, Berkeley, Berkeley, CA 94720, USA

## Abstract

mTORC1 regulates cell growth and catabolism in response to fluctuations in nutrients through phosphorylation of key substrates. The tumor suppressor FLCN is a RagC GTPase activating protein (GAP) that regulates mTORC1 phosphorylation of TFEB, controlling lysosome biogenesis and autophagy. Here, we determined the cryo-EM structure of the active FLCN complex (AFC) containing FLCN, FNIP2, the N-terminal tail of SLC38A9, the RagA^GDP^:RagC^GDP.BeFx-^ GTPase dimer, and the Ragulator scaffold. Relative to the inactive lysosomal FLCN complex (LFC) structure, FLCN reorients by 90°, breaks its contacts with RagA, and makes new contacts with RagC that position its Arg164 finger for catalysis. Disruption of the AFC-specific interfaces of FLCN and FNIP2 with RagC eliminated GAP activity in vitro and led to nuclear retention of TFE3, with no effect on mTORC1 phosphorylation of S6K or 4E-BP1. The structure thus provides a roadmap to discover TFEB-selective mTORC1 antagonists.

**One-Sentence Summary:** The cryo-EM structure of the active FLCN RagC GAP complex provides a structural basis for TFEB/TFE3 substrate-selective targeting of mTORC1.

## Main Text

The mechanistic target of rapamycin complex 1 (mTORC1) plays a central role in the response to fluctuations in nutrients, growth factors, and energy in cells by altering the balance between cell growth and catabolism. In nutrient-rich environments, mTORC1 activates anabolic processes including protein and lipid synthesis while inhibiting catabolic ones such as autophagy (*1, 2*). mTORC1 regulates its pro-anabolic programs through phosphorylation of key targets, some of which contain a conserved TOS motif (*3, 4*). The microphthalmia family (MiTF/TFE) of transcription factors, including transcription factors EB and E3 (TFEB/TFE3), are pro-catabolic mTORC1 substrates that activate autophagy and lysosome biogenesis (*5-7*). Among well-known mTORC1 substrates, TFEB/TFE3 lack the TOS motif responsible for docking some other substrates onto mTORC1 via its Raptor subunit. Also, in contrast to other mTORC1 substrates, TFEB/TFE3 phosphorylation and negative regulation are completely dependent on the activity of the RagC GAP, FLCN (*8, 9*).

The Rag-Ragulator complex consists of two Rag guanosine triphosphatases (commonly referred to as Rags) that function as obligate heterodimers and are anchored to the lysosomal membrane by the pentameric Ragulator (*10, 11*). The Rags respond to lysosomal and cytoplasmic nutrient levels, which regulate the proportion of inactive (RagA or BGDP:RagC or DGTP) and active (RagA or BGTP:RagC or DGDP) Rag dimers (*12*). Conversion between active and inactive states is regulated by the RagA/B-specific GAP, GATOR1, and by the RagC/D-specific GAP, folliculin: folliculin interacting protein 2 (FLCN:FNIP2) (*13, 14*). In response to amino acid levels, as sensed by the transceptor SLC38A9 among other mechanisms (*15-17*), FLCN stimulates the conversion of Rags to their active form enabling mTORC1 recruitment and interaction with TFEB/TFE3. Upon phosphorylation, TFEB/TFE3 binds to 14-3-3 proteins and is sequestered in the cytosol where it remains inactive. The regulation of TOS motif-containing mTORC1 substrates (*3*), including regulators of mRNA translation and cell proliferation, such as 4E-BP1 and S6K1 is unaffected (*9, 18*). Taken together, FLCN directly and selectively controls mTORC1 regulation of TFEB/TFE3 but no other substrates (*19-21*). These findings support the existence of separate mechanisms for control of mTORC1-dependent anabolic versus catabolic programs, and highlight a therapeutic potential for FLCN to be targeted as a means of mTOR inhibition for treating LSDs and neurodegenerative diseases, with fewer toxic effects on cell proliferation and translation.

The transient nature of the GAP interaction with Rag-Ragulator has proven challenging for visualizing details necessary for deducing the mechanism of GAP activity. Previous structural investigation into FLCN interaction with Rag-Ragulator revealed a stable inhibited complex, the lysosomal folliculin complex (LFC) (*9, 22*). This begged the question as to how FLCN binds to RagC to promote nucleotide hydrolysis, and what are the interfaces in FLCN essential for stimulating catalysis? We sought to understand how these residues interact with the RagC nucleotide binding pocket, and what makes FLCN specific for RagC. We previously found that the amino acid transporter, SLC38A9, promotes FLCN GAP activity by inserting its cytosolic domain in the cleft between the RagA and C G-domains and thereby disrupting the LFC (*16*). Here, we trapped FLCN in its GAP-competent binding mode using a fusion construct with the N-terminal of SLC38A9 (SLC38A9^NT^), determined the cryo-EM structure, and deduced the mechanism of FLCN GAP activity essential for interrogating this interaction during drug development.

To provide a structural framework for understanding the mechanism of FLCN GAP activity, we stabilized the transient complex and determined the cryo-EM structure, hereafter referred to as the Active Folliculin Complex (AFC). We began by generating a stable complex of FLCN:FNIP2, Rags, Ragulator and SLC38A9^NT^, utilizing a mutant FLCN^F118D^ (hereafter referred to as FLCN) known to disrupt the LFC without affecting GAP activity (*9*). To tether FLCN to the Rag-Ragulator complex, we designed a fusion construct containing a 10-residue glycine-serine linker between the C-terminus of SLC38A9^NT^ and the N-terminus of FNIP2 (Fig. 1A). The C-terminal tail of SLC38A9^NT^ contains ∼20 disordered residues, effectively increasing the linker length to ∼30 residues allowing for up to 90 Å in length. We used a phosphate analog (BeF_3_) during complex formation to capture FLCN bound to Rag-Ragulator (reference on BeF_3_ use). Rags were loaded into a state mimicking starvation conditions (RagA^GDP^:RagC^GTP^) and incubated with Ragulator to form a stable complex. Next, FLCN and the SLC38A9^NT^-FNIP2 (SLC-FNIP2) fusion protein were added to RagA^GDP^:RagC^GTP^ – Ragulator immediately followed by addition of BeF_3_ (Fig. S1). Incubating the complex overnight, we generated a nucleotide ground-state analog that trapped FLCN in an active conformation and allowed for isolation of a stable active FLCN complex via size exclusion chromatography (Fig. 1B, 1C).

**Figure 1.**
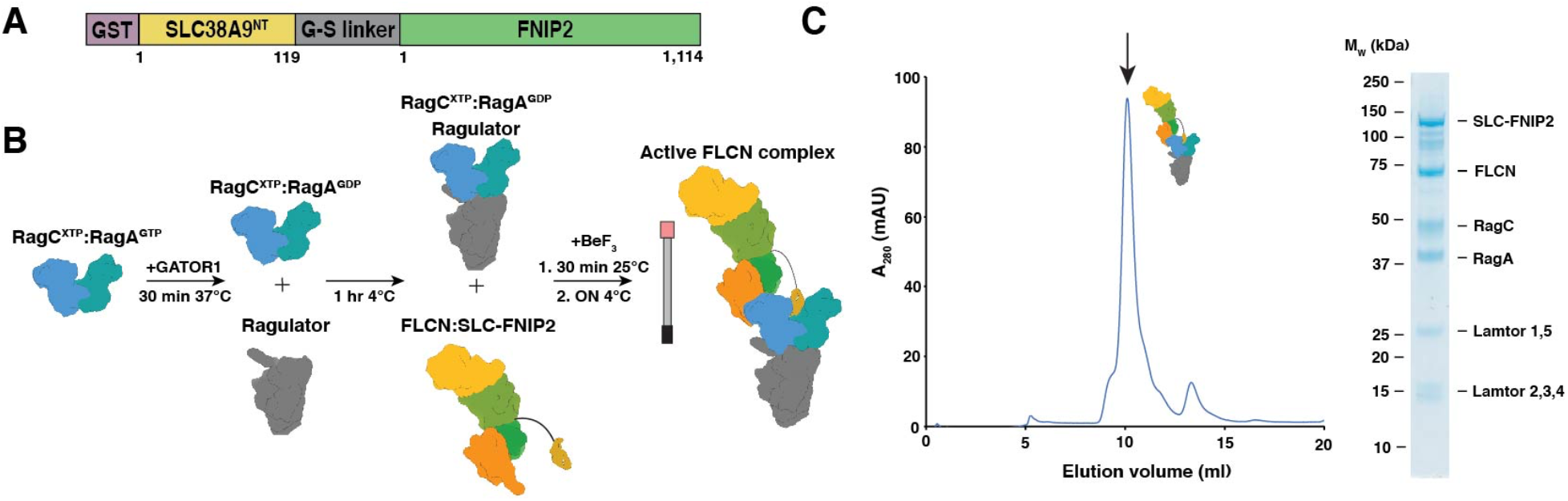
Active FLCN complex formation. (A) Construct design for SLC38A9^NT^-FNIP2 fusion construct. (B) Complex formation workflow. (C) Size exclusion profile for active FLCN complex. Fraction containing all components of complex is labelled with a black arrow and visualized on adjacent gel.

The cryo-EM structure of the AFC was determined to an overall resolution of 3.5 Å (Table 1, Fig. S2). The cryo-EM density was of sufficient quality to model the ordered mass of the complex constituting the AFC (Fig. S3 and S4). In the AFC, the longin domains of FLCN and FNIP2 interact with the RagC G-domain such that the FLCN:FNIP2 approaches Rag-Ragulator from the side (Fig. 2B, 2C). The SLC38A9^NT^ sits in the cleft between the Rags and does not interact directly with FLCN:FNIP2, suggesting it does not play a role in positioning the GAP. The overall dimensions of the complex are ∼120 Å by ∼180 Å (Fig. 2C). To interact with RagC in a GAP-competent conformation, FLCN:FNIP2 reorients 90 degrees and transitions from its elongated position in the LFC to bind the side of RagC in the AFC (Movie S1). In the LFC, FLCN:FNIP2 interacts with both RagC and RagA extensively, with 870 Å^2^ and 645 Å^2^ interfaces, respectively. In contrast, in the AFC conformation, FLCN:FNIP2 interacts solely with RagC, burying 1,141 Å^2^ of surface area with RagC, but none with RagA.

**Figure 2.**
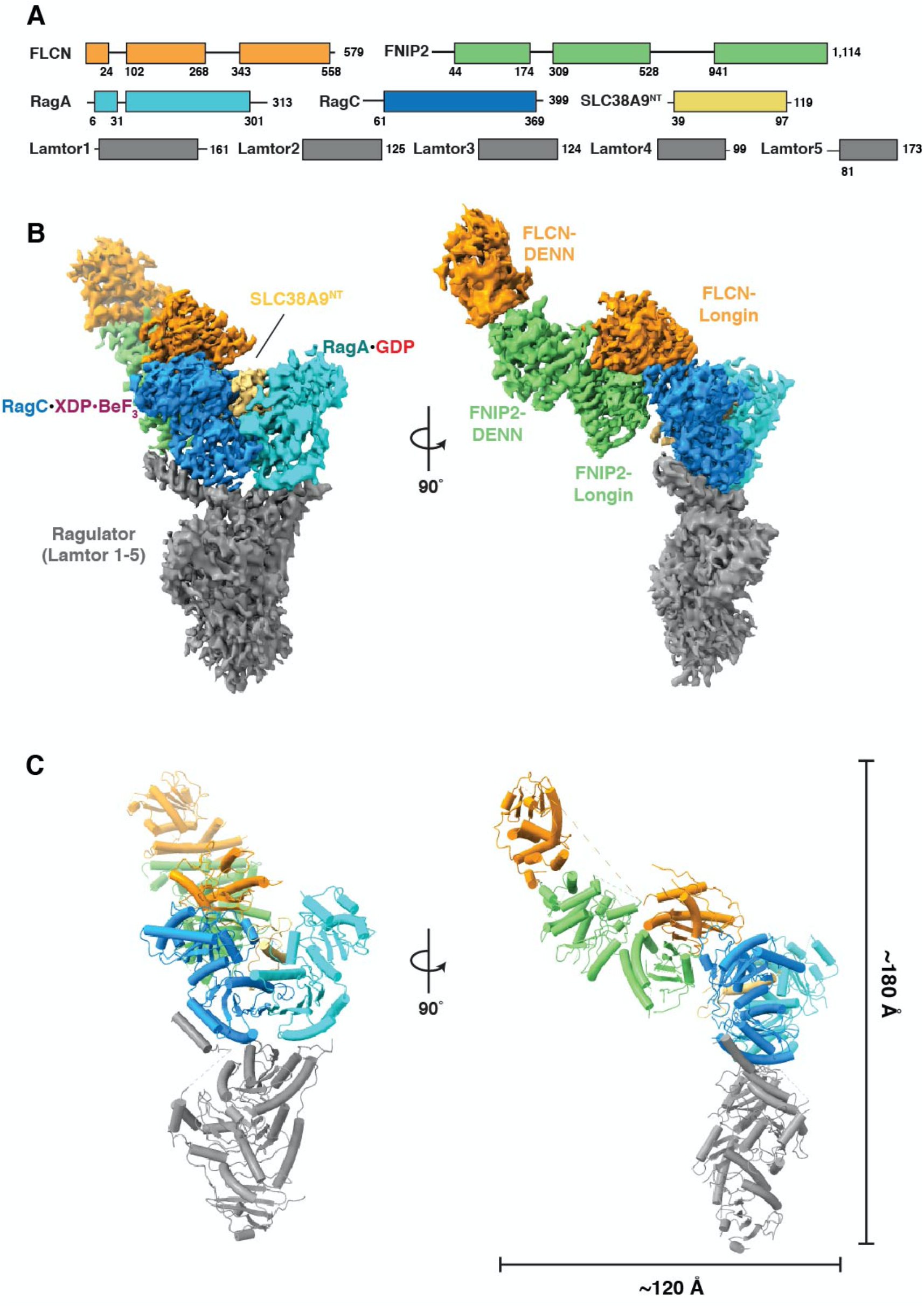
Cryo-EM structure of active FLCN complex. (A) Domain organization for components of active FLCN complex. (B) Composite map for complex. Focused map of interface was combined with overall map to produce composite map containing highest resolution information for each component. (C) Reconstructed model for AFC with over dimensions of the side view of the complex.

To visualize the AFC catalytic complex in as much detail as possible, we used local refinement at the FLCN-RagC interface to increase the resolution to 3.16 Å and allow for clear placement of the nucleotide analog and previously identified arginine finger (Arg-164) in RagC nucleotide-binding domain (NBD) (*9*) (Fig. S2 and S3D). As bound to GDP-BeF^-^_3_, RagC adopts an architecture analogous to GTP-bound RagC and Gtr2 (*23, 24*) (Fig. 3A). In this state, the switch I region is ordered and forms a lid at the top of the binding pocket. Additional density in the RagC NBD, adjacent to the BeF^-^_3_, aligned with the placement of the highly conserved Arg-164. The position of FLCN Arg-164 resembled the geometry arginine fingers in previously identified arginine fingers in Rab1 GAP structures trapped with BeF^-^_3_ nucleotide analogs (*25, 26*) (Fig. S5). The nucleotide binding pocket is composed of primarily charged residues with a 534 Å ^2^ surface area, while the overall volume of the larger surrounding pocket is 865 Å^3^ (*27*) (Fig. 3B).

**Figure 3.**
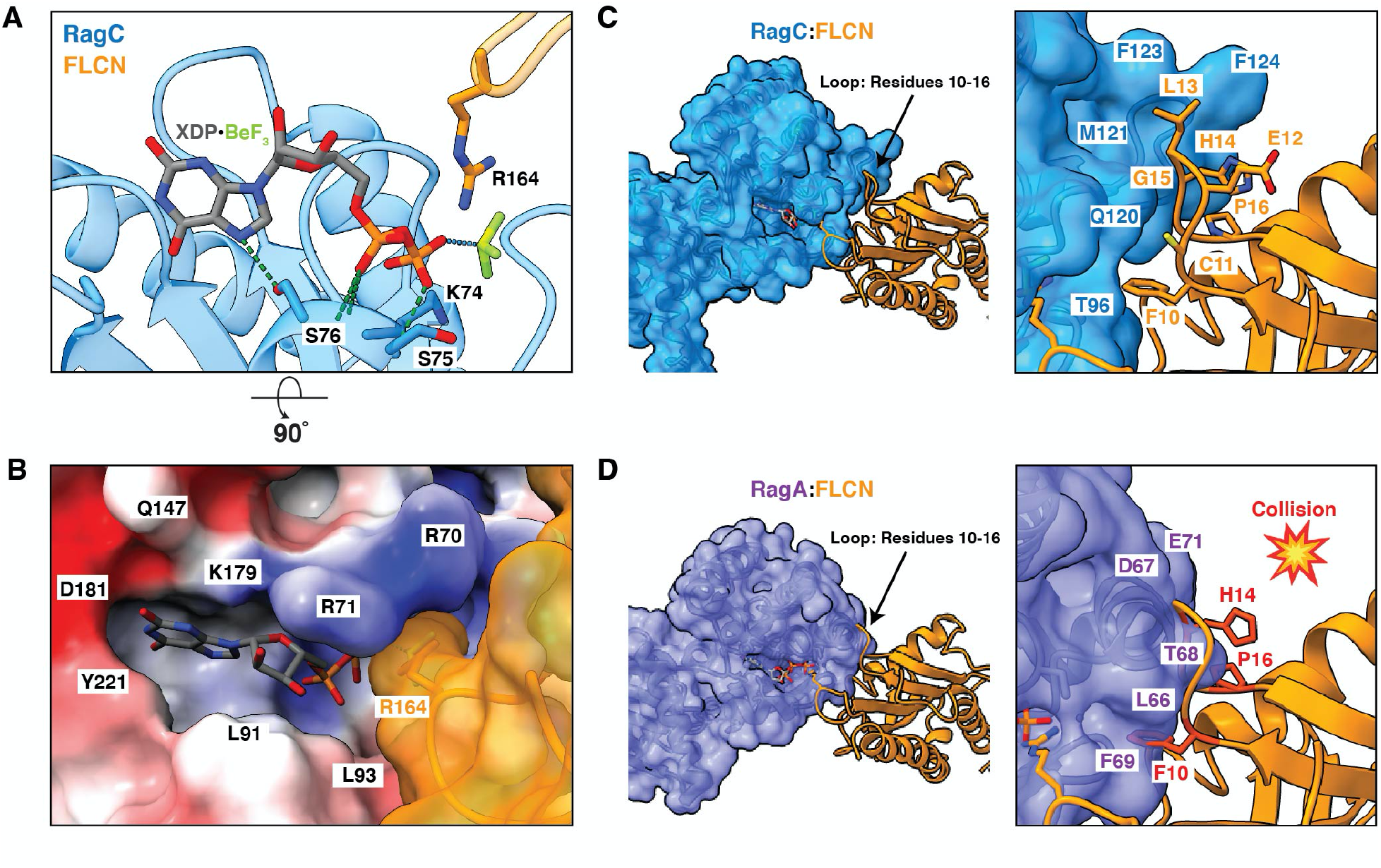
Structure of the active FLCN-RagC GAP interface. (A) Nucleotide binding pocket of RagC (blue) in AFC containing XDP-BeF_3_ nucleotide analog. Location of FLCN (orange) arg-finger indicated. Hydrogen bonds between nucleotide and RagC residues (blue sticks) in surrounding pocket represented by dashed lines (green). (B) Surface representation of RagC binding site with FLCN arg-finger (orange). RagC colored based on electrostatic potential. (C) RagC (blue) FLCN (orange) interface in AFC. Loop residues 10-16 in FLCN indicated as sticks (black labels). (D) RagA (purple) FLCN interface modelled with GTP-bound RagA (PDB:6S6D). Residues in FLCN loop that clash with RagA are shown in red.

To determine the basis for FLCN specificity for RagC over RagA, we modeled the GAP interaction with RagA-GTP. The FLCN loop containing residues 10-16 was accommodated when interacting with RagC (Fig. 3C). In contrast, in the RagA model, FLCN (Phe^16^, His^14^, Pro^16^) clashed with RagA at the interface (Fig. 3D). We sought to further explore FLCN specificity for RagC by understanding why GATOR1 does not function as a GAP for RagC. We modelled RagC with GATOR1 and observed residues in GATOR1 (Thr^18^ and Pro^19^) that clashed with RagC at the interface (Fig. S6B). We also compared the RagC structure to that of Arf1, which is a substrate of another longin domain GAP, SMCR8 (*28, 29*). In the Arf1 model, FLCN (Leu^13^ and His^14^) clash with Arf1 residues (Fig. S6A), rationalizing specificity for RagC. To determine why other longin domain GAPs do not function to stimulate nucleotide hydrolysis of RagC, we modelled RagC with SMCR8, a longin GAP that acts on Arf GTPases. The binding between SMCR8 and RagC is incompatible due to clashes between SMCR8 (Glu^62^ and Phe^63^) and RagC (Fig. S6C).

We analyzed the interface whereby FLCN:FNIP2 interacts with RagC. Two hydrophobic residues FLCN Phe10 and FNIP2 Val146 form prominent contacts in the RagC binding interface interacting with Thr96, Lys78, Met121 and Phe128 (Fig. 4A). We hypothesized that these residues were critical for GAP activity. To validate the importance of these residues, we created two mutants FLCN^F10D^ (F10D:Phe^10^→Asp) and FNIP2^V146E^ (V146E:Val^146^→Glu). We used a tryptophan-based GAP activity assay to monitor the activity of wild-type (WT) FLCN:FNIP2 and mutants FLCN^F10D^:FNIP2 and FLCN:FNIP2^V146E^ toward inactive Rags (RagA^GDP^:RagC^GTP^) (*9, 16*). The GAP activity of the FLCN:SLC-FNIP2 fusion protein was identical to the FLCN:FNIP2 without the SLC38A9 N-terminus fusion (Fig. S7). Both FLCN^F10D^:FNIP2 and FLCN:FNIP2^V146E^ abolished GAP activity as compared to WT, FLCN:FNIP2 confirming their importance in facilitating GAP binding (Fig. 4).

**Figure 4.**
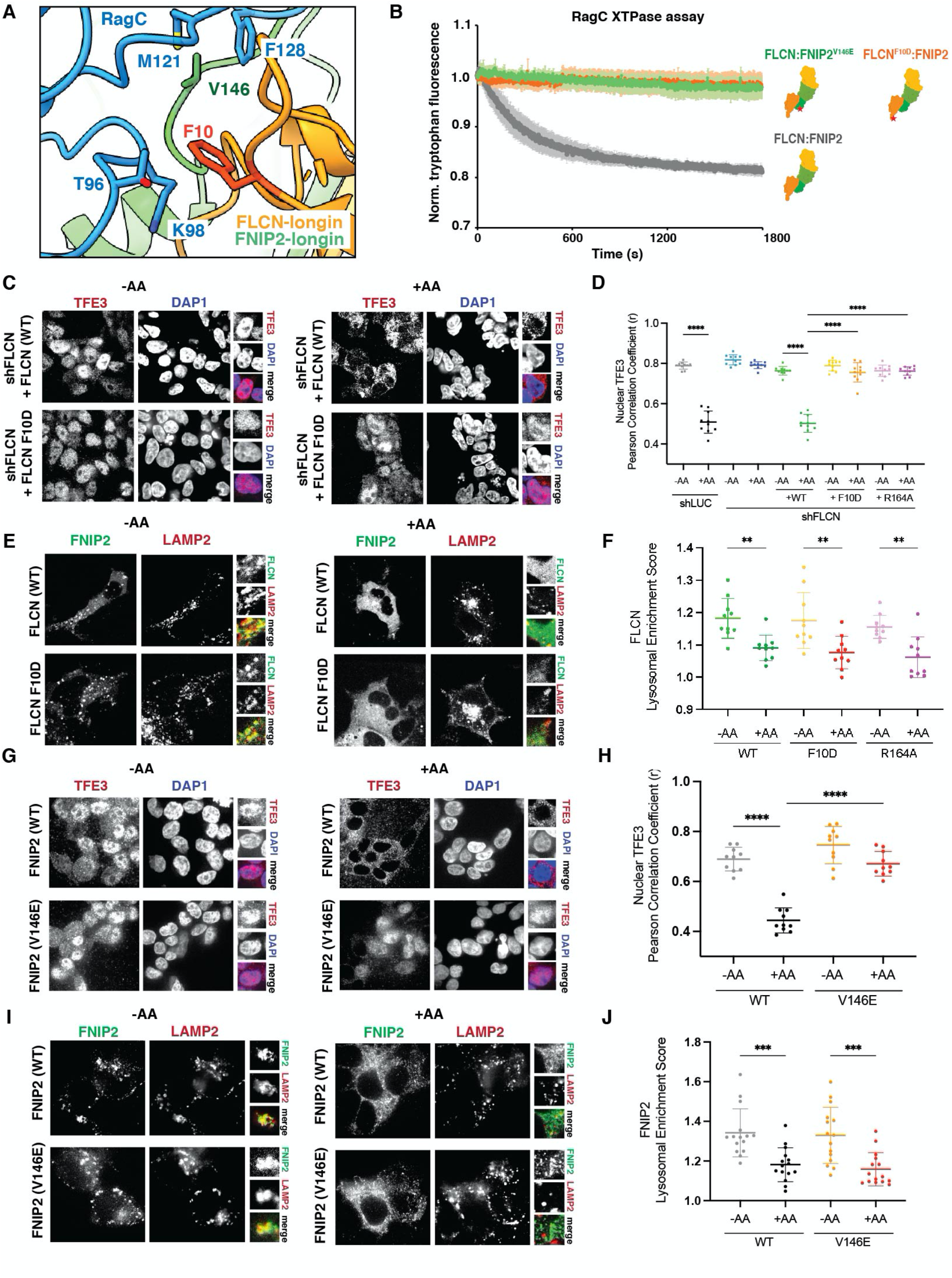
Hydrophobic residues in the FLCN:RagC interface necessary for GAP activity. (A) RagC (blue) and FLCN (orange):FNIP2 (green) interface in AFC. Position of FLCN:FNIP2 residues mutated at interface indicated. (B) Tryptophan fluorescence XTPase assay with FLCN: FNIP2 mutants. FLCN^WT^: FNIP2 (gray), FLCN^F10D^: FNIP2 (orange), or FLCN: FNIP2^V146E^ (green) was incubated with RagA^GDP^:RagC^XTP^. Plotted are means ± SEM. n=3 replicates. (C) Immunofluorescence images of human embryonic kidney 293T (HEK293T) cells stably expressing the indicated genes shRNA and FLCN rescue constructs. Cells were starved for 2 hours for amino acids (-AA) or restimulated with complete DMEM for 2 hours (+AA). (D) Quantification of TFE3 nuclear localization for immunofluorescence images and the positive control FLCN^164^ identified previously (*9*) and shown in Fig. S8 immunofluorescence images (means ± SD, n = 10 fields for all conditions). **** p< 0.0001. (E) Immunofluorescence images of human embryonic kidney 293A (HEK293A) cells expressing the indicated FLCN constructs. Cells were starved for 1 hour for amino acids (-AA) or restimulated with amino acids for 15 min (+AA). (F) Quantification of FLCN lysosomal enrichment score for immunofluorescence images (means ± SD, n = 15 cells for all conditions) ** p< 0.01. (G) Immunofluorescence images of HEK293T cells stably expressing the FNIP2^WT^ or FNIP2^V146E^ constructs. Cells were starved for 2 hours for amino acids (-AA) or maintained in complete media (+AA). (H) Quantification of TFE3 nuclear localization for immunofluorescence images (means ± SD, n=10 fields for all conditions) **** p< 0.0001. (I) Immunofluorescence images of human embryonic kidney 293T (HEK293T) cells stably expressing FNIP2:V5 construct. LAMP2 was used as lysosomal marker. Cells were starved for 2 hours for amino acids (-AA) or maintained in complete media (+AA). (J) Quantification of FNIP2 lysosomal enrichment score for immunofluorescence images (means ± SD, n = 15 cells for all conditions) *** p< 0.001.

We sought to investigate whether these GAP-inactive mutants, FLCN^F10D^ and FNIP2^V146E^, affected the ability of Rag-Ragulator to activate mTORC1 with respect to its various substrates. We used cell-based assays to image the localization of MiT/TFE family transcription factors TFE3/TFEB. In amino-acid deprived cells reconstituted with either WT FLCN or FNIP2, both FLCN and FNIP2 were localized to the lysosome (Fig. 4G) and TFE3/TFEB was present in the nucleus (Fig. 4C and 4G). Cells expressing FLCN^F10D^ and FNIP2^V146E^ during starvation resulted in lysosomal localization and nuclear localization of TFEB (Fig. 4C, 4E, and 4G). When cells were restimulated with amino acids, those reconstituted with WT FLCN and FNIP2 resulted in translocation of TFE3 to the cytoplasm. In contrast, in cells expressing FLCN^F10D^ and FNIP2^V146E^ TFE3 remained nuclear, phenocopying the catalytic loss-of-function mutant FLCN^R164^ (*9*) (Fig. S8) and was not sequestered to the cytoplasm despite high cellular nutrients (Fig. 4C, 4E). These results further supported that the mutants FLCN^F10D^ and FNIP2^V146E^ eliminated GAP activity and confirmed that loss of GAP activity resulted in mis-regulation of mTORC1 and TFEB/TFE3. Immunoblot analysis confirmed that the mutants did not have any effect on the phosphorylation of the TOS-motif containing substrates S6K and 4E-BP1 (Fig. S9).

The cryo-EM structure presented in this work reveals that FLCN-FNIP2 undergoes a dramatic re-orientation to convert from the inactive LFC to the active AFC. The AFC structure confirmed the prediction that FLCN activates nucleotide hydrolysis of RagC through the catalytic arginine finger Arg-164 (*9, 30*). We identified features on the FLCN surface that are complementary to RagC and confer the high degree of specificity that is seen biologically. Our structure revealed that the loop containing residues 10-16 in FLCN make it incompatible as a GAP for other small GTPases. In the AFC structure, SLC38A9^NT^ sits in the cleft between the Rags, while FLCN:FNIP2 binds only RagC. This confirms the previously proposed model that SLC38A9^NT^ displaces inactive FLCN:FNIP2 in the LFC to allow for binding in the GAP competent mode (*16*). SLC38A9^NT^ does not, however, play a role in GAP activity through interaction with FLCN or either Rag nucleotide binding pocket in the AFC, as further supported by the near-identical RagC-GAP activity of FLCN:FNIP2 with or without the SLC38A9^NT^ fusion.

RagA is regulated by a different GAP complex, GATOR1. A recent cryo-EM structure of the GAP-competent interaction between Rag-Ragulator and GATOR1 suggested GATOR1 is stabilized by both Rag GTPases in its GAP mode (*31*). The overall geometry and the role of the Arg finger in the active GATOR1 complex resembles that of FLCN-FNIP2 seen here, as well as that of another longin-domain based GAP, the C9orf72-SMCR8 complex with ARF1 (*28*). In comparison to GATOR1, the structure of the AFC revealed that FLCN solely interacts with RagC to stimulate GTP hydrolysis. This finding is consistent with observations that while GATOR1 RagA GAP activity depends on the RagC nucleotide state, the RagC GAP activity of the AFC is independent of the RagA nucleotide state (*16*).

The past few years have seen a paradigm shift in understanding how the MiTF family of transcription factors is regulated downstream of mTORC1 in a uniquely FLCN and RagC^GDP^-dependent manner (*8, 9, 18, 19, 21, 32, 33*). The new knowledge of FLCN structural mechanism provided by the AFC structure can potentially be exploited for therapeutic benefit. The AFC presents itself as an attractive avenue for the long-standing goal of developing substrate selective mTORC1 inhibitors which can upregulate autophagy and lysosomal capacity without toxic consequences for immune function, epithelial renewal and other physiological processes that require mTORC1-dependent anabolic programs. Such inhibitors could be highly specific for TFEB/E3 upregulation, in turn promoting autophagy and increasing lysosomal capacity. The latter could be of potential benefit for treating LSDs, while both could potentially benefit neurodegenerative diseases involving intracellular protein aggregates. The structural insights into the AFC presented here fulfill a key step necessary to leverage FLCN as a drug target and develop selective mTORC1 inhibitors for TFEB/TFE3 upregulation.

## Acknowledgements

We thank Zhicheng Cui and Aaron Joiner for discussions, Rick Hooy for workstation support and Dan Toso and Jonathan Remis for electron microscope support.

## Funding

This work was supported by the NIH grants R01 GM111730 (J.H.H.) and R01 GM130995 (R.Z.), a National Science Foundation Graduate Research Fellowship (R.M.J.), and an Italian Cancer Research Association (AIRC) postdoctoral fellowship (R.P.).

## Author contributions

R.M.J. and J.H.H. conceived and designed research, R.M.J. and R.P. carried out research, A.L.Y. and S.F. trained R.M.J. in cryo-EM, R.Z. and J.H.H. supervised research, R.M.J. and J.H.H. wrote the first draft, and all authors edited the manuscript.

## Competing interests

J.H.H. is a co-founder and shareholder of Casma Therapeutics and receives research funding from Casma Therapeutics, Genentech, and Hoffmann-La Roche. R.Z. is a cofounder and shareholder of Frontier Medicines and receives research funding from Genentech.

## Data and materials availability

Coordinates and density are being deposited in the RCSB and EMDB, respectively. The fusion construct used to generate the AFC is being deposited in Addgene.org.

## Supplementary Materials

### Materials and Methods

#### Cloning and Protein Purification

The pCAG-GST-SLC38A9^NT^-FNIP2 fusion construct was designed to contain a ten-residue glycine-serine linker. Codon-optimized DNA coding for SLC38A9^NT^ and the C-terminal linker was amplified by polymerase chain reaction (PCR). DNA containing SLC38A9^NT^ and the C-terminal linker was subcloned into the pCAG-GST vector containing codon-optimized DNA coding for FNIP2. The pCAG-GST-FNIP2 vector was linearized using KpnI and XhoI restriction sites. DNA containing SLC38A9^NT^ and C-terminal linker was inserted N-terminal to FNIP2 by Gibson assembly. HEK293-GNTI cells were transfected with 1mg DNA at 2:1 ratio (FLCN: SLC38A9^NT^-FNIP2) and 3mg P.E.I per 1L of cells at 2E6 cells/ml. Cells were pelleted at 2000 xg for 20 min at 4 °C and purified according to *Lawrence et al. (9*).

Mutants FLCN F10D and FNIP2 V146E were generated by site-directed mutagenesis using KAPA HiFi HotStart ReadyMix (Roche) from pCAG-FLCN and pCAG-GST-FNIP2 respectively. HEK293-GNTI cells were transfected with 1mg DNA at 2:1 ratio (FLCN:FNIP2) 3mg P.E.I per 1L of cells at 2E6 cells/ml. Cells were pelleted at 2000 xg for 20 min at 4 °C and purified according to *Lawrence et al. (9*).

RagA:RagC^D181N^ and Ragulator were expressed and purified from Sf9 cells via baculovirus infection as described in *Lawrence et al*. (*9*). GATOR1 was expressed and purified from HEK293-GNTI cells as described in *Lawrence et al. (9*).

#### Nucleotide Loading

Purified RagA:RagC^D181N^ heterodimers were diluted 1:10 (v/v) in 1X phosphate-buffered saline (PBS) containing 5mM EDTA and 0.5 mM TCEP for 10 minutes at 25°C. Desired tri-phosphate nucleotides were added at a 10-fold molar excess over Rags and incubated for 30 minutes at 25°C. Next, MgCl2 was added to a final concentration of 20 mM and incubated for 10 minutes at 25°C. Excess nucleotides were removed using a PD-10 desalting column (Cytiva) and wash buffer containing 25 mM HEPES, 130 mM NaCl, 2.5 mM MgCl2, 2 mM EGTA, pH 7.4 and 0.5mM EDTA. To obtain the inactive state for Rag heterodimers, RagA^GTP^:RagC^XTP^ was incubated with GATOR1 at a 1:100 GAP:Rag molar ratio at 37 °C for 30 minutes.

#### Active FLCN Complex Assembly

To assemble the active FLCN complex, RagA:RagC^D181N^ heterodimers in desired nucleotide state were incubated for 1 hour at 25°C with a 1.2x molar excess of Ragulator. FLCN: SLC38A9^NT^-FNIP2 was added to Rag-Ragulator complex at a 2-fold molar excess followed immediately by the addition of 0.5 M BeF_3_. BeF_3_ was prepared by mixing BeSO_4_ and NaF to final concentration of 0.5mM and 5mM respectively and incubating for 20 minutes at 25°C prior to addition to complex. The complex containing Rags, Ragulator, FLCN: SLC38A9^NT^-FNIP2 and BeF_3_ was incubated at 25°C for 30 minutes, then at 4°C for 16 hours. Following incubation, the complex was centrifuged at 17,000 xg for 10 minutes and then loaded onto a Superose 6 (GE Healthcare) column equilibrated in wash buffer containing 25 mM HEPES, 130 mM NaCl, 2.5 mM MgCl2, 2 mM EGTA, pH 7.4 and 0.5mM EDTA. Fractions were analyzed via SDS PAGE and those containing assembled active FLCN complex were collected, concentrated to 1 mg/ml and immediately frozen onto grids for imaging via cryo-EM.

#### Cryo-EM Grid Preparation and Imaging

For data collection of active FLCN complex, 3 μl sample (1 mg/ml protein in 25 mM HEPES, 130 mM NaCl, 2.5 mM MgCl2, 2 mM EGTA, 0.5 mM TCEP, pH 7.4) was deposited onto freshly glow-discharged (PELCO easiGlow, 45 s in air at 20 mA and 0.4 mbar) holey carbon grids (C-flat: 2/1-3Cu-T). FEI Virobot Mark IV was used to blot grids for 2 seconds with a blot force of 20 (Whatman 597 filter paper) at 4 °C and 90-100 % humidity and subsequentially plunged into liquid ethane. The Titan Krios G3i microscope equipped with a Gatan Quantum energy filter (slit width 20 eV) and a K3 summit camera at a defocus of -1.0 to -2.0 μm was used to record 4968 movies. Automated image acquisition was performed using SerialEM (*34*) recording four movies per 2 μm hole with image shift. Image parameters are summarized in Table S1.

#### Cryo-EM Data Processing

The data processing workflow is summarized in Figure S1. In short, Relion-3.1.1 wrapped MotionCor2 program was used to motion-correct gain-corrected movies (*35, 36*). Motion-corrected micrographs were imported into cryosparc2 v3.3.1 (*37*). Patch CTF estimated (multi) was used for CTF determination and two rounds of cryosparc2 blob picker with a diameter range of 150Å-220Å was used to generate 1,165,732 and 1,025,735 particles. Particles were extracted with a box size of 440×440 pixels in cryosparc2. A series of 2D classifications followed by an ab-initio-reconstruction was used to generate three reference maps. The resulting 3D maps were used for consecutive rounds of heterogenous refinement from the initial particle sets following a round of 2D classification to remove obvious ‘junk’. The resulting particle sets from round 1 and round 2 of picks were merged and duplicates were removed. The combined particle set contained 177,018 particles. A final round of homogenous refinement resulted in a 3.47Å map at 0.143 FSC. A mask was generated surrounding the interface between RagC and FLCN:FNIP2 using UCSF Chimera and imported into cryosparc2 v3.3.1 where it was lowpass filtered and dilated (*38*). The mask was used for subsequent local refinement and resulted in a focused map reaching a resolution of 3.16Å.

#### Atomic Model Building and Refinement

The coordinates from RagC were based on GTP-bound crystal structure (3LLU), the coordinates for FLCN and FNIP2 were based off the LFC (6NZD) and all five Ragulator, RagA and SLC38A9^NT^ were based on Rag-Ragulator-SLC38A9^NT^ structure (6WJ2). All coordinates were rigid body fitted separately into the density map using UCSF Chimera. The interface was refined using a map generated from local refinement separately by iterative rounds of Phenix real-space refinement and manual correction in coot (*39*). Secondary structure restrains were enabled during real-space refinement. The interface was combined with the remaining components of the complex in coot and refined using iterative round of Phenix real-space refinement and ISOLDE (*40*). The final model was validated using MolProbity (*41*). Coordinate models can be found in the Protein Data Bank (PDB) with accession code XXXX.

#### Tryptophan Fluorescence RagC XTPase Assay

Tryptophan fluorescence experiments were performed in triplicate according to *Fromm et al*. (*16*).

#### Cell Culturing

HEK293T cells were grown in Dulbecco’s modified Essential Media (DMEM, Thermo Fisher Scientific) supplemented with 10% (v/v) fetal bovine serum (FBS, VWR), 100 U/ml penicillin and 100 U/ml streptomycin (Life Technologies) and were maintained at 37 °C and 5% CO_2_.

#### Reagents and Antibodies

The following antibodies were employed in this study: phospho-T389 S6K1 (9234S), S6K1 (2708S), phospho-S65 4EBP (9451S), 4EBP (9644S and 9452S), FLCN (3697), TFE3 (14779), TFEB (37785), V5 (13202), FLAG (14793) from Cell Signaling Technology; LAMP2 (sc-18822) from Santa Cruz Biotechnology.

#### RNAi

The control pLKO.1-LUC shRNA vector and the pLKO.1-FLCN lentiviral shRNA vector were obtained respectively from Addgene (Plasmid #30324) and the RNAi Consortium (TRCN0000237886). Target sequences are:

shLUC: TCCTAAGGTTAAGTCGCCCTCG.

shFLCN: GATGGAGAAGCTCGCTGATTT

RNAi experiments were prepared as described below.

#### Lentiviral Infection and Stable Cell Line Generation

Lentivirus was generated using the PEI method. 5 μg of a lentiviral vector (desired construct in pLJM1-PURO) was combined with 3.75 μg of psPAX2 and 1.25 μg pCMG.2 viral packaging plasmids in 500 μl optiMEM (Thermofisher), mixed, and then combined with 60 μl 1 mg/mL P.E.I. This solution was incubated at room temperature for 30 minutes, then added to a 10 cm dish containing 2 million recently plated 293T cells.

24-48 hours after that, media containing virus was collected from cells, spun down at 1,300 x rpm for 5 minutes to remove cell debris, and sterile filtered. Cells were plated at a density of 200,000-300,000 cells per well in a 6-well plate in the presence of Polybrene transfection reagent (Sigma-Aldrich) and infected with 10% of the total virus from one 10 cm plate of the indicated constructs. 24 hours later, media was supplemented with 200 nM puromycin.

#### Live Cells Treatments

HEK293T cells were seeded and allowed to attach overnight in complete DMEM medium. 24h after seeding, cells were rinsed and incubated in amino acid-free RPMI for 2 hours (-AA) and restimulated with amino acids for 15 minutes or overnight with complete media (+AA). Amino acid solutions were prepared from powders and the final concentration of amino acids in the media was the same as in commercial RPMI. Torin1 treatment was performed for 4 hours in complete media.

#### Western Blotting

HEK-293T cells were plated in a 6-well plate at 1,000,000 cells/well in complete media. After the treatments (see above), the medium was removed and 150 µl of Lysis Buffer (150 mM NaCl, 20 mM HEPES [pH 7.4], 2 mM EDTA, 1% Triton X-100, and one tablet of EDTA-free protease inhibitors per 50 ml) were added to each well. Total extracts were collected from each well after scraping, rotated over-end for 10 minutes, and then centrifuged at 13,000 × g for 10 min at 4 °C. Samples were normalized to a total concentration of 1 mg/mL protein and combined with protein sample buffer, then boiled for 5 minutes at 95 °C.

Ten micrograms of protein were loaded into each lane of Tris-Glycine precast gels (Thermo Fisher Scientific). PVDF membranes were blocked in 5% skim milk dissolved in Tris-buffered saline with Tween 20 for 1 hour at RT. Membranes were incubated overnight at 4 °C with the indicated primary antibodies, all diluted in 5% skim milk prepared in TBST. Secondary antibodies conjugated to horseradish peroxidase were used for protein detection. β-Actin was used as loading controls.

#### Immunofluorescence

HEK-293T cells were plated on fibronectin-coated glass coverslips in 12-well plates, at 300,000 cells/well. The following day, cells were subjected to amino acid depletion/or complete media restimulation (see above) and then fixed in 4% paraformaldehyde (in PBS) for 15 min at RT. The coverslips were rinsed twice with PBS and cells were permeabilized with 0.1% (w/v) saponin in PBS for 10 min. After rinsing twice with PBS, the slides were incubated overnight at 4°C with the indicated primary antibody in 5% normal donkey serum, then rinsed with PBS, and incubated with fluorophore-conjugated secondary antibodies produced in goat or donkey (Life Technologies, diluted 1:500 in 5% normal donkey serum) for 45 min at room temperature in the dark. Coverslips were then washed three times in PBS and mounted on glass slides using Vectashield Antifade Mounting Medium (Vector Laboratories) containing DAPI stain. All images were collected on a Nikon Ti-E inverted microscope (Nikon Instruments, Melville, NY) equipped with a Plan Apo 60x oil objective. Images were acquired using a Zyla 5.5 sCMOS camera (Andor Technology), using iQ3 acquisition software (Andor Technology).

#### Quantitation of Lysosomal Enrichment Score

For immunofluorescence datasets, a home-built MATLAB (Mathworks) script was used to determine the lysosomal enrichment of both V5-FNIP2 and FLAG-FLCN stains as previously described in *Lawrence et al*. (*9*). Briefly, a single cell was manually selected in the lamp channel and its nucleus was excluded from further analysis. Then, a mask was created in the LAMP2 channel to segment cellular pixels into LAMP2 (lysosomal) or non-LAMP2 (cytosolic) pixels. This mask was then applied to the non-LAMP2 channel. The Lysosomal Enrichment Score was determined by dividing the average intensity of pixels in the lysosomal region by the average intensity of the pixels in the cytosolic region. For each condition, at least twenty cells were analyzed from different multi-channel images.

#### Quantitation of Cytoplasmic:Nuclear TFE3 Ratio

To measure TFE3 nuclear localization, the Pearson’s coefficient of at least 10 different fields was measured using the ImageJ plugin JACoP (*42*).

**Fig. S1.**
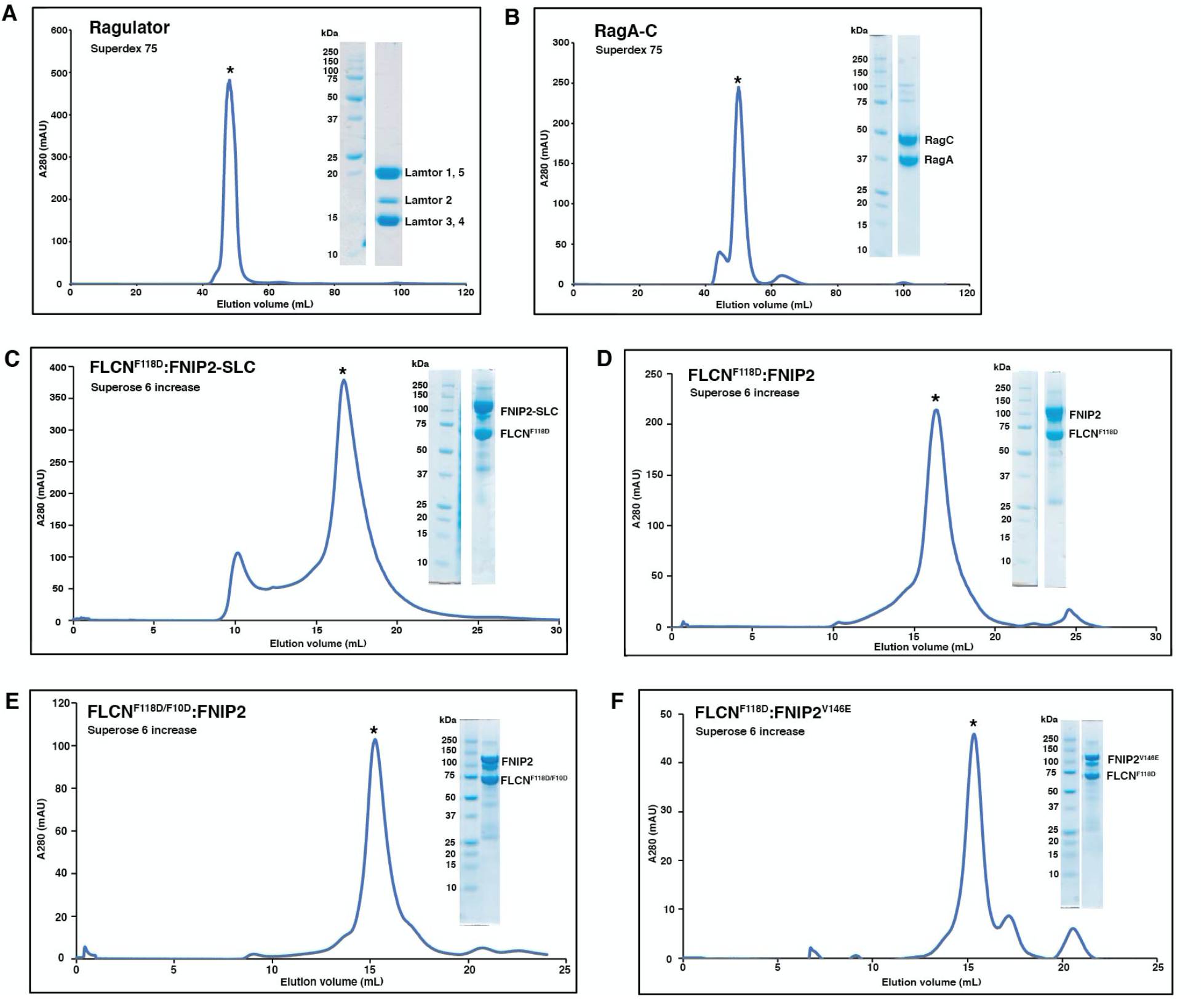
Purification of individual active FLCN components. (A) Superdex 75 SEC elution profile for Ragulator and Coomassie blue stained SDS-PAGE analysis of protein sample collected from peak indicated with asterisk. (B) Superdex 75 SEC elution profile for Rag GTPases and Coomassie blue stained SDS-PAGE analysis of protein sample collected from peak indicated with asterisk. (C) Superose 6 increase SEC elution profile for FLCN^F118D^:FNIP2-SLC fusion and Coomassie blue stained SDS-PAGE analysis of protein sample collected from peak indicated with asterisk. (D) Superose 6 increase SEC elution profile for FLCN^F118D^:FNIP2 and Coomassie blue stained SDS-PAGE analysis of protein sample collected from peak indicated with asterisk. (E) Superose 6 increase SEC elution profile for FLCN^F118D/F10D^:FNIP2 and Coomassie blue stained SDS-PAGE analysis of protein sample collected from peak indicated with asterisk. (F) Superose 6 increase SEC elution profile for FLCN^F118D^:FNIP2^V146E^ and Coomassie blue stained SDS-PAGE analysis of protein sample collected from peak indicated with asterisk.

**Fig. S2.**
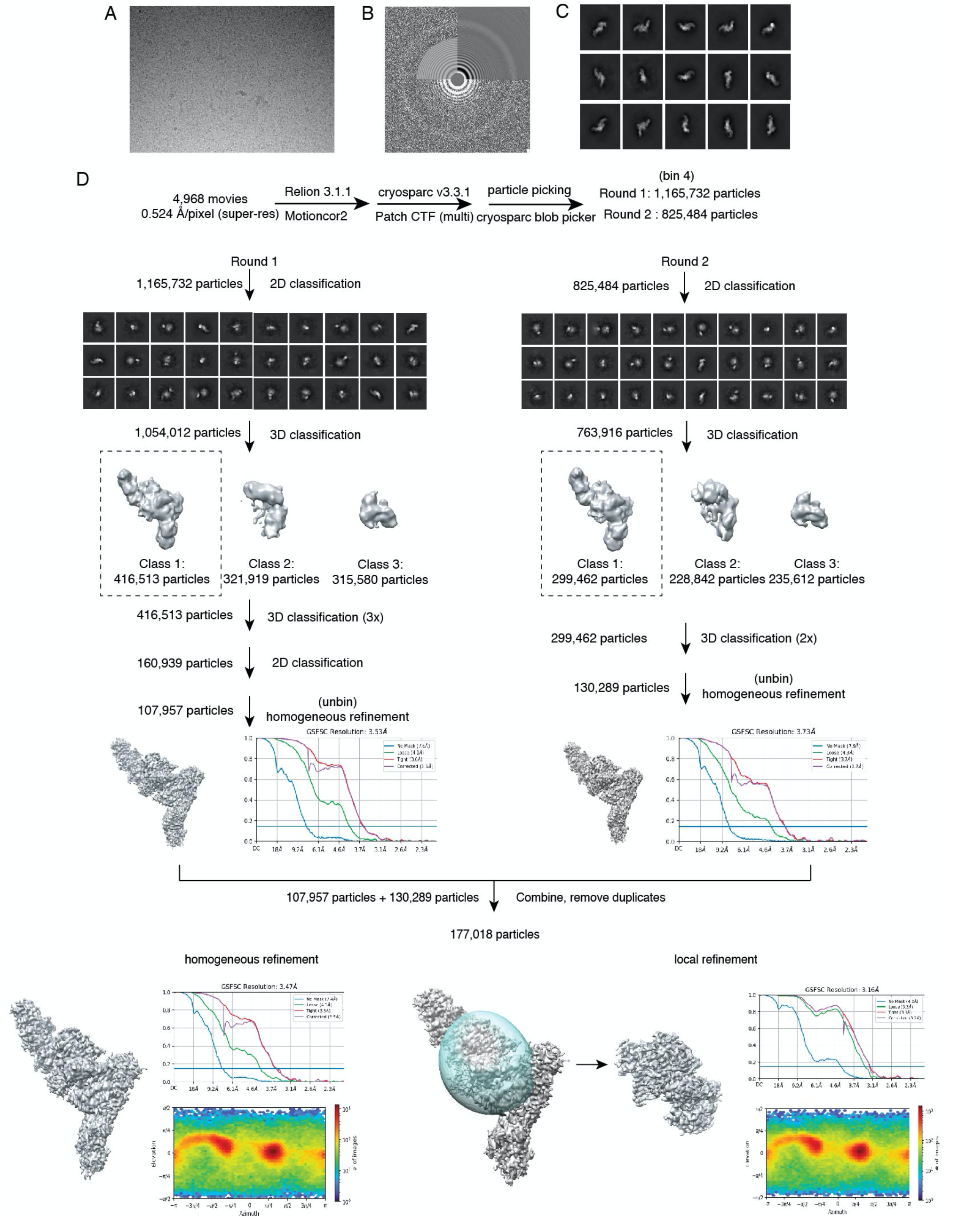
Cryo-EM data processing pipeline. (A) Representative cryo-EM micrograph (B) Power spectrum and CTF estimation of micrograph shown in (A). Exemplary 2D class averages for the active FLCN complex. (D) Data processing pipeline for final map determination.

**Fig. S3.**
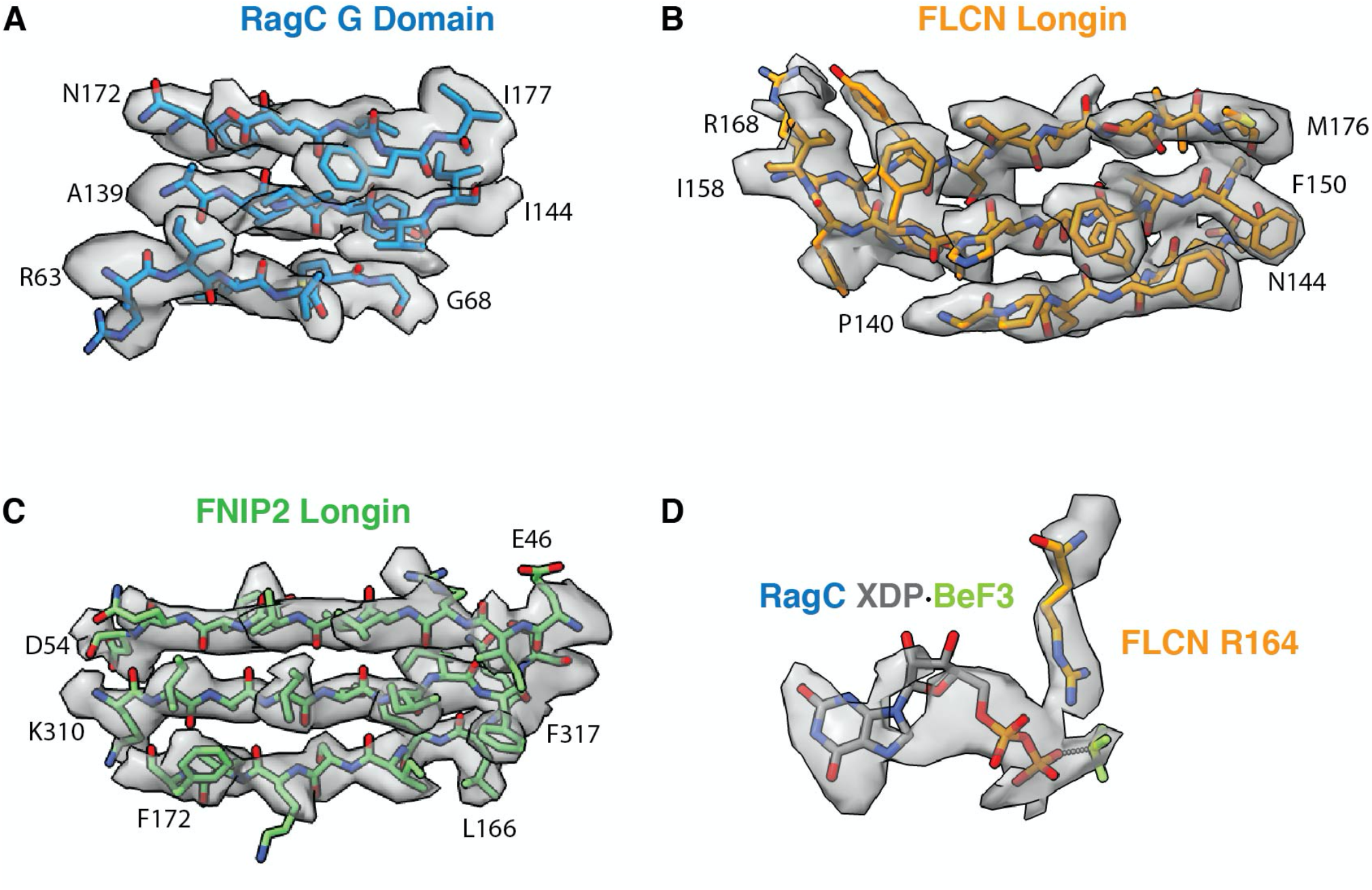
AFC map-model fit. Representative refined coordinate model fit in cryo-EM density for (A) RagC (B) FLCN (C) FNIP2 (D) RagC nucleotide and FLCN arginine finger.

**Fig. S4.**
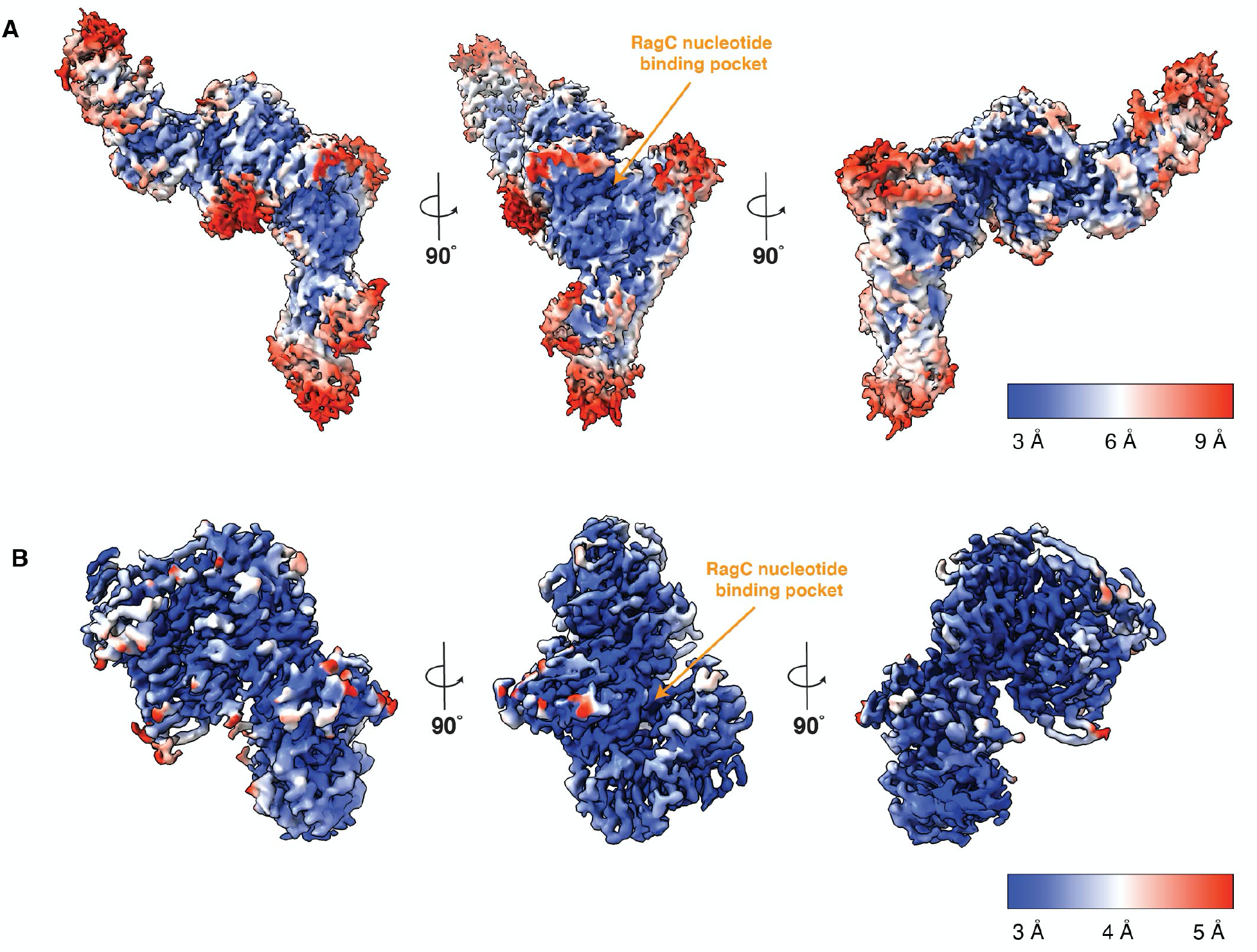
AFC local resolution estimation. (A) Overall cryo-EM map for AFC colored according to local resolution. Resolution ranges from 3 Å to 9 Å. (B) Cryo-EM map of RagC – FLCN^login^:FNIP2^longin^ interface generated from focused refinement colored according to local resolution. Resolution ranges from 3 Å to 6 Å.

**Fig. S5.**
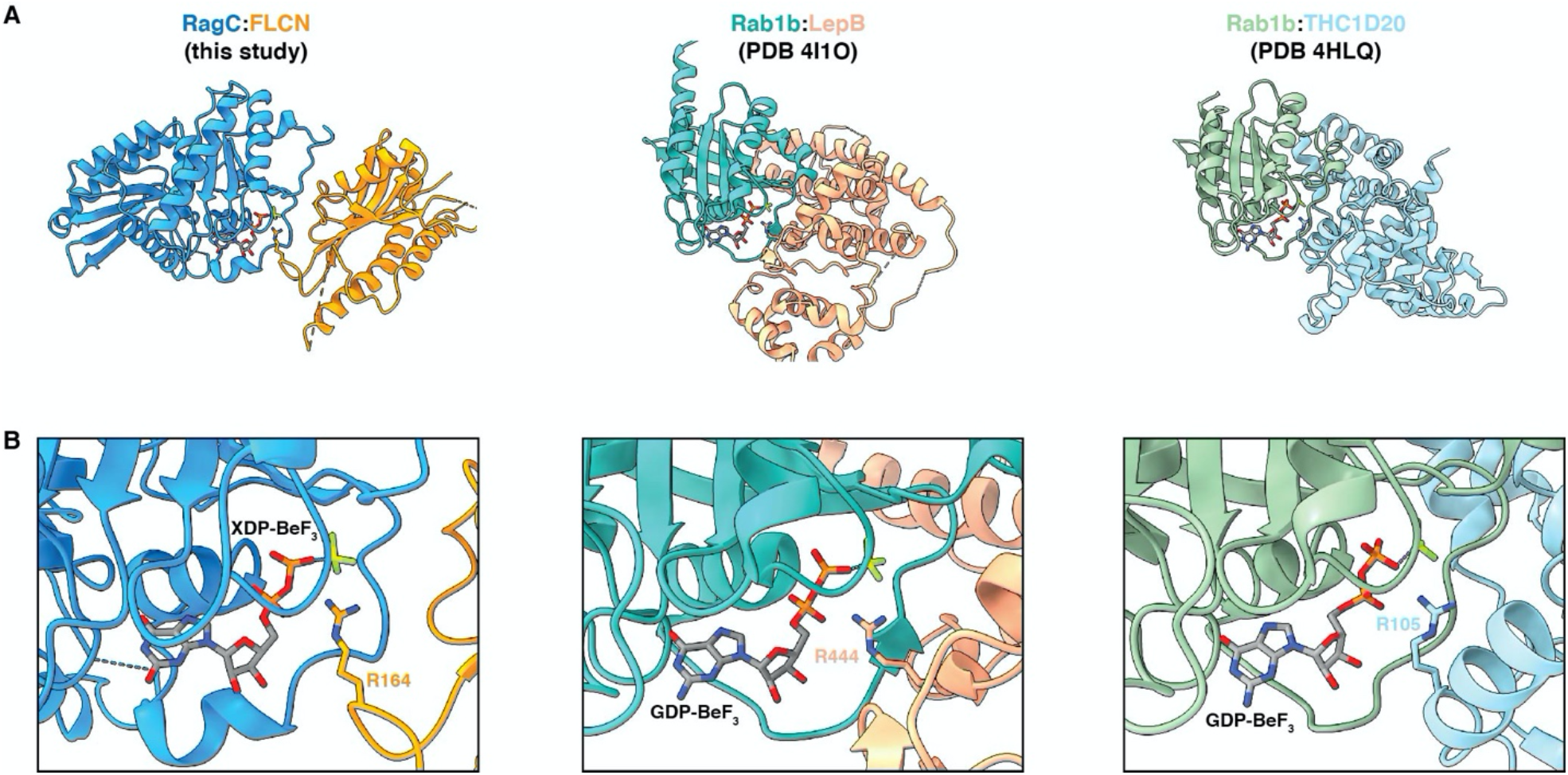
Structural comparison between RagC:FLCN and other small GTPases:GAP. (A) Rab1b:LepB and Rab1b:THC1D20. (A) Structural comparison of RagC:FLCN (AFC structure), Rab1b:LepB (PDB 4I1O) and Rab1b:THC1D20 (PDB 4HLQ). Structures are shown in the same orientation. (B) Close-up of the nucleotide binding pocket with catalytic arginine-finger residue positioned in active site.

**Fig. S6.**
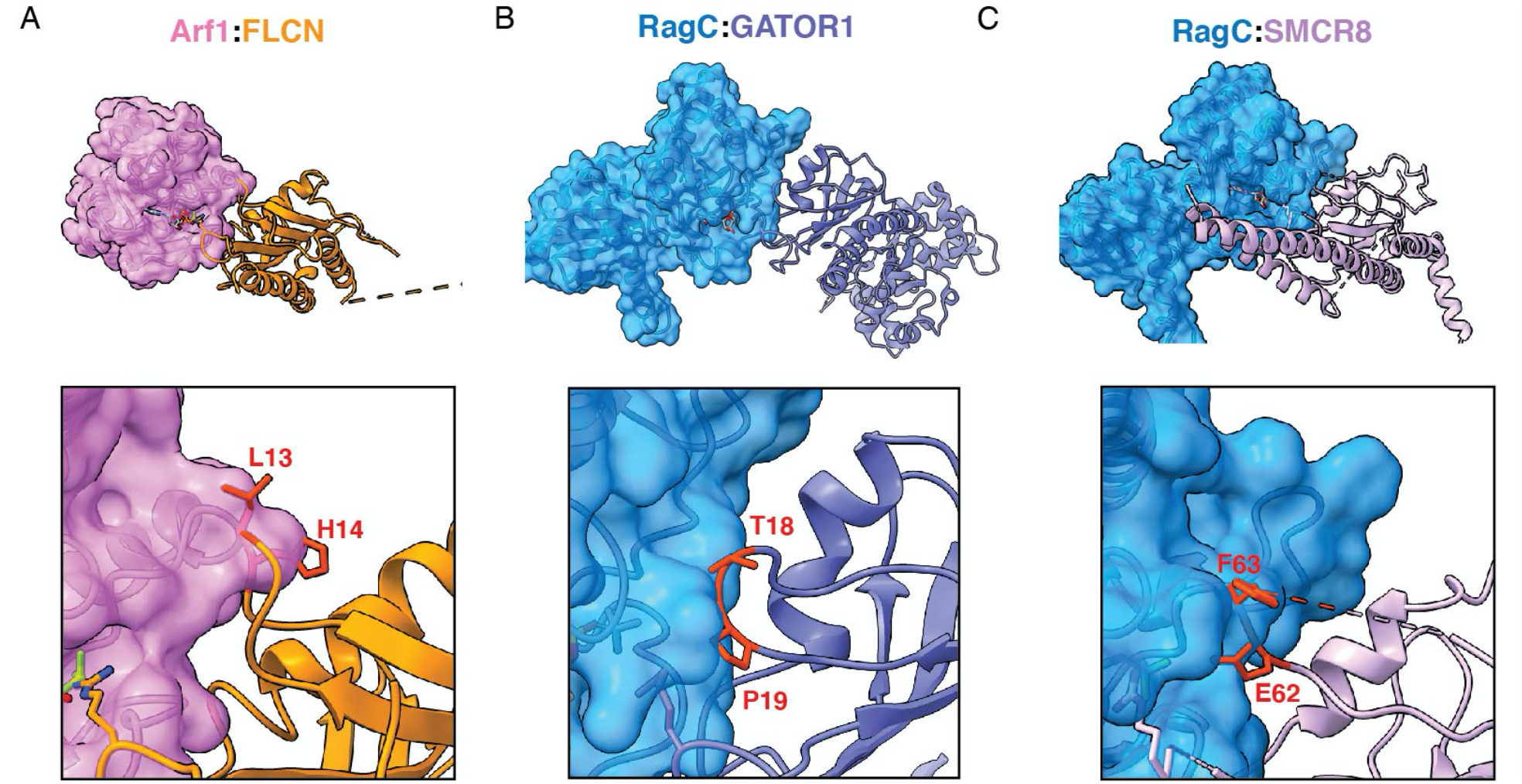
Structural investigation of RagC:SMCR8 and Arf1:FLCN interface. (A) Model of Arf1 (pink) and FLCN (orange) interface generated by aligning Arf1 from Arf1:SMCR8 structure (PDB: 7MGE) with RagC in AFC structure (*27*). Residues in FLCN at interface that clash are indicated and labelled in red. (B) Model of RagC (pink) and SMCR8 (orange) interface generated by aligning SMCR8 from Arf1:SMCR8 structure (PDB: 7MGE) with FLCN in AFC structure (*27*). Residues in SMCR8 at interface that clash are indicated and labelled in red.

**Fig. S7.**
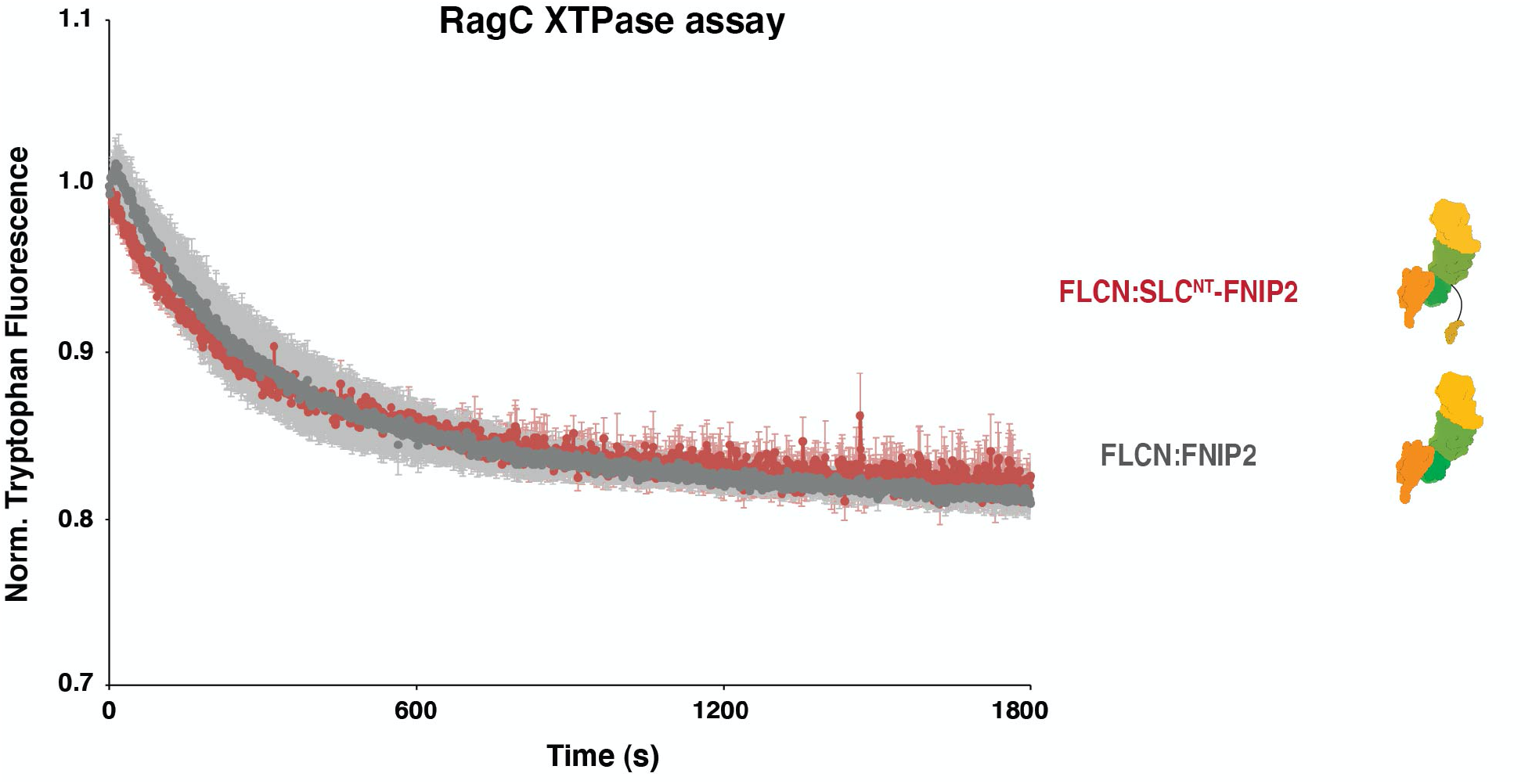
GAP activity of FLCN:SLC-FNIP2. Tryptophan fluorescence XTPase assay with FLCN:SLC-FNIP2 fusion construct. FLCN^F118D^: FNIP2 (gray), FLCN^F118D^: SLC-FNIP2 (red), was incubated with RagA^GDP^:RagC^XTP^. Plotted are means ± SEM. n=3 replicates.

**Fig. S8.**
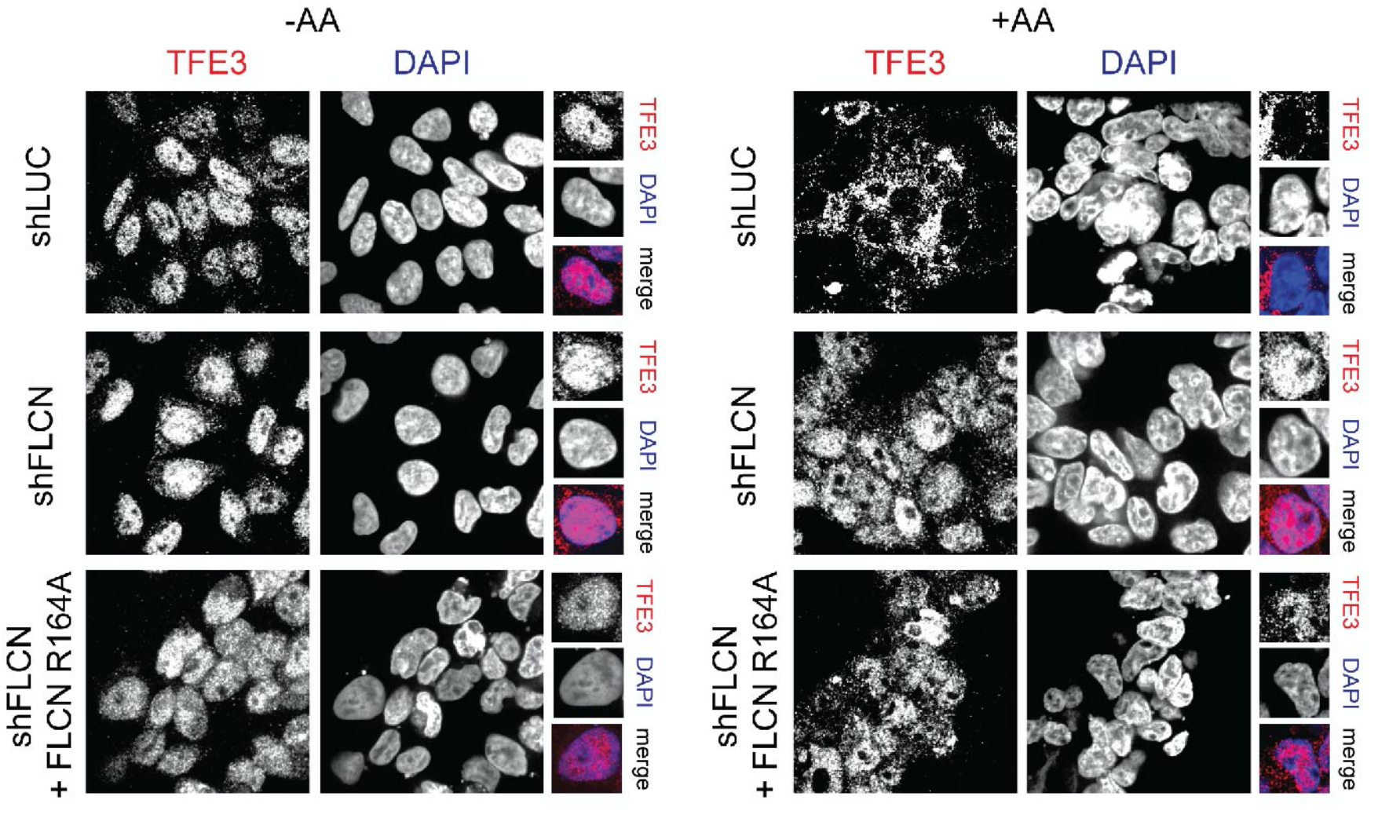
Immunofluorescence for FLCN^R164^. Immunofluorescence images of human embryonic kidney 293T (HEK293T) cells stably expressing the indicated genes shRNA and FLCN rescue constructs. Cells were starved for 2 hours for amino acids (-AA) or restimulated with complete DMEM for 2 hours (+AA).

**Fig. S9.**
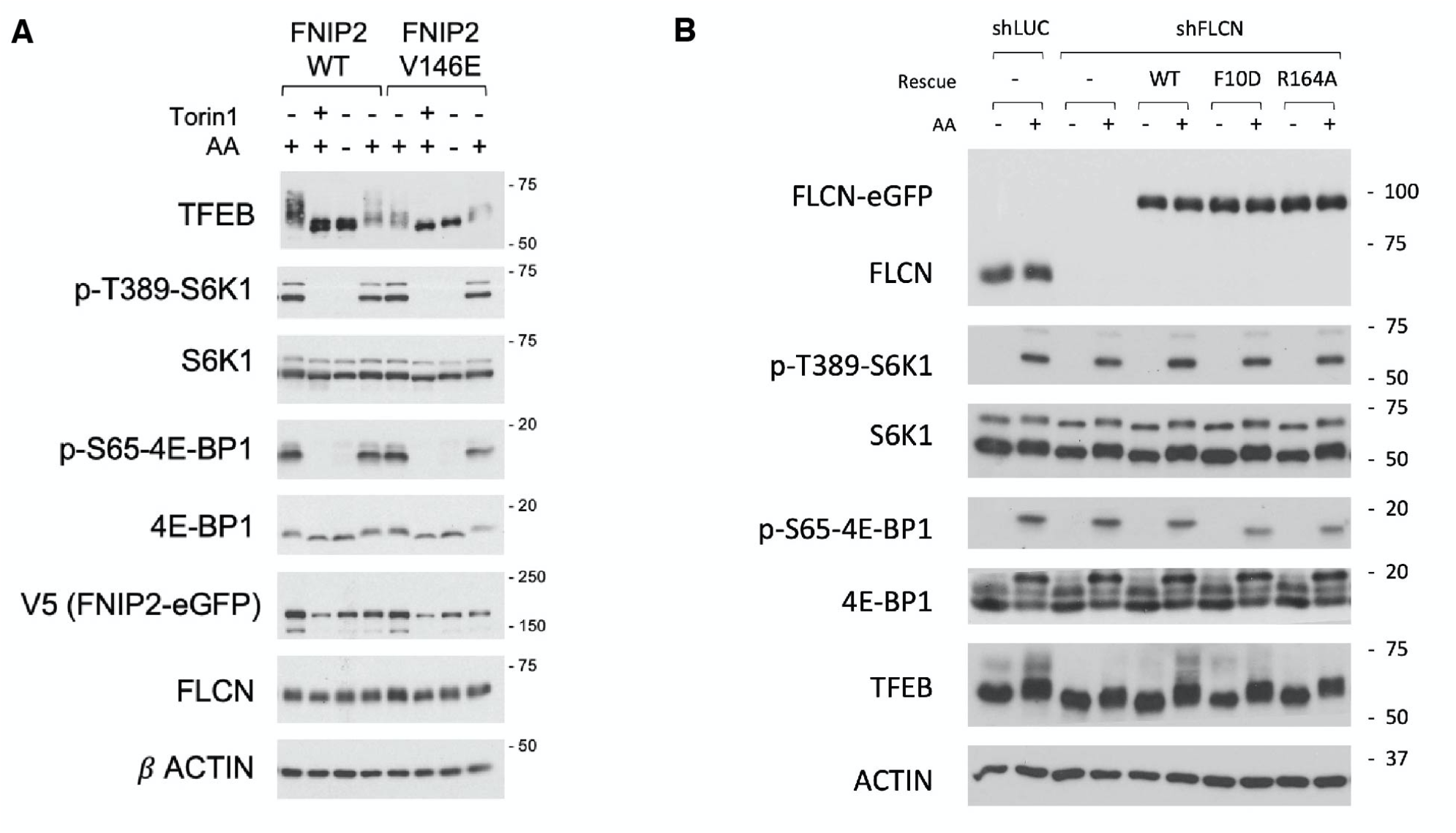
Immunoblot analysis for mTORC1 targets. (A) HEK-293T cells stably expressing FNIP2^WT^ or FNIP2^V146E^ were either untreated or treated with Torin1, or starved of amino acids for 2 hours, or starved and then restimulated with amino acids for 15 minutes. Cells were lysed, followed by immunoblotting for the indicated proteins and phospho-proteins. (B) HEK-293T cells stably expressing shRNAs targeting the indicated genes were starved of amino acids for 2 hours, or starved and then restimulated with amino acids for 15 minutes. Cells were lysed, followed by immunoblotting for the indicated proteins and phospho-proteins.

**Table S1.** Cryo-EM data acquisition and image processing.

|  | Active FLCN complex |
| --- | --- |
| **Data acquisition** |  |
| Microscope | Titan Krios |
| Voltage (kV) | 300 |
| Camera | Quantum-K3 Summit |
| Magnification | 165,000 |
| Pixel size (Å) | 0.524 (super-resolution) |
| Cumulative exposure (e<sup>-</sup>/Å<sup>2</sup>) | 50 |
| Energy filter slit width (eV) | 20 eV |
| Defocus range (µm) | -1.0 to -2.0 |
| Automation software | SerialEM |
| Exposure navigation | Image Shift |
| Number of movies | 4968 |
| **Image processing** |  |
| Initial picked particles (no.) | Round 1: 1,165,732 Round 2: 825,484 |
| Final refined particles (no.) | 177,018 |
| Map resolution (Å) | Overall: 3.53 Interface: 3.16 |
| FSC threshold | 0.143 |

**Table S2.**
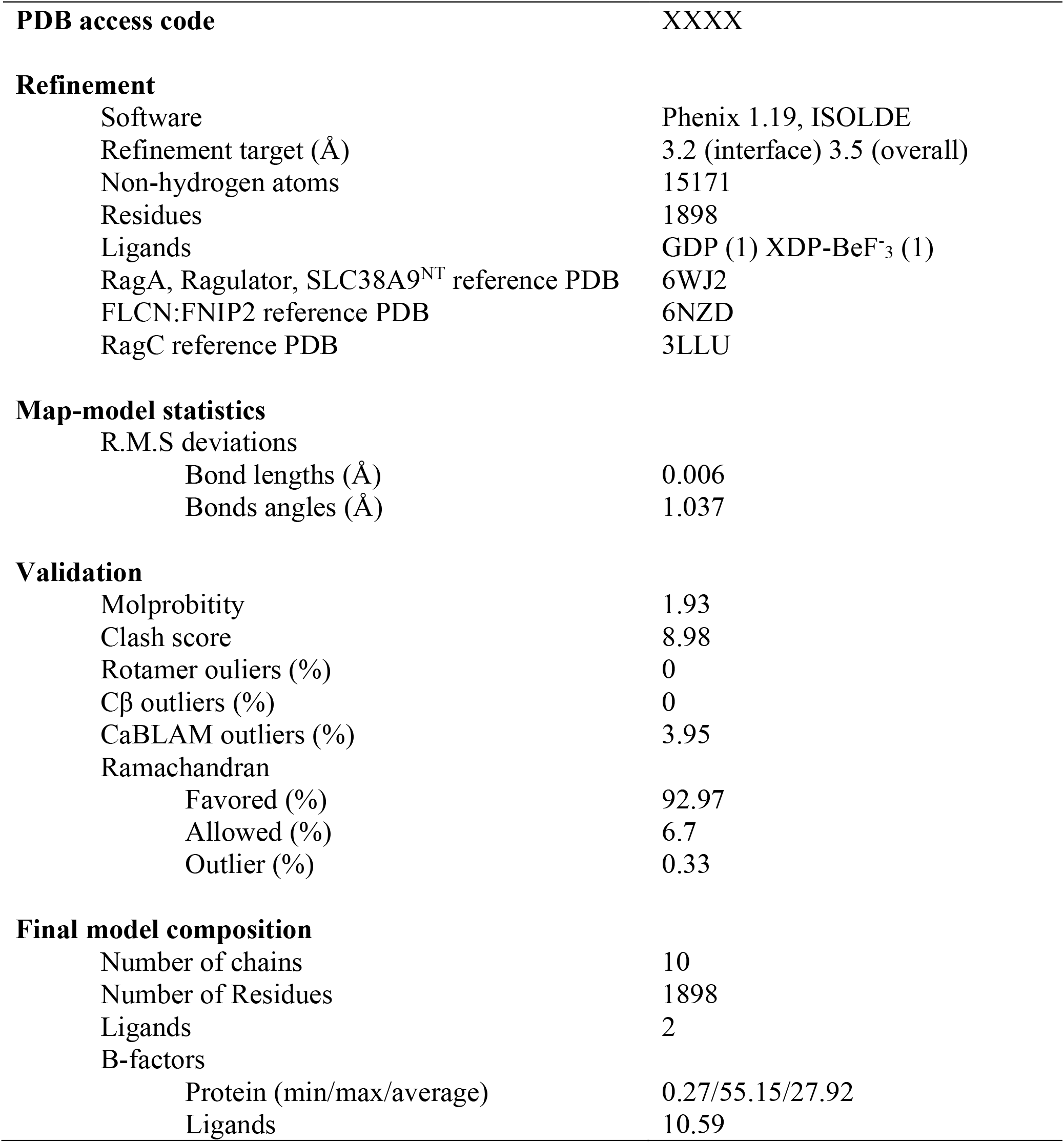
AFC coordinate model refinement and assembly.

**Movie S1. Structural rearrangement of FLCN:FNIP2 between LFC and AFC**. Visualization of the inactive (LFC) and active (AFC) binding modes of FLCN:FNIP2 to Rag-Ragualtor.

## Notes

### Summary of Updates

This version corresponds to the initial submission to the journal which has provisionally accepted the associated manuscript and complies with the journal's embargo policy.

